# I-KCKT allows dissection-free RNA profiling of adult *Drosophila* intestinal progenitor cells

**DOI:** 10.1101/2020.06.27.175174

**Authors:** Kasun Buddika, Jingjing Xu, Ishara S. Ariyapala, Nicholas S. Sokol

## Abstract

The adult Drosophila intestinal epithelium is a model system for stem cell biology, but its utility is limited by current biochemical methods that lack cell type resolution. Here, we describe a new proximity-based profiling method that relies upon a GAL4 driver, termed *intestinal-kickout-GAL4* (*I-KCKT-GAL4*), exclusively expressed in intestinal progenitor cells. This method used UV cross-linked whole animal frozen powder as its starting material to immunoprecipitate the RNA cargoes of transgenic epitope-tagged RNA binding proteins driven by *I-KCKT-GAL4*. When applied to the general mRNA-binder, poly(A)-binding protein, the RNA profile obtained by this method identified 98.8% of transcripts found after progenitor cell sorting, and had low background noise despite being derived from whole animal lysate. We also mapped the targets of the more selective RNA binder, Fragile Mental Retardation Protein, using enhanced CLIP, and report for the first time its binding motif in Drosophila cells. This method will therefore enable the RNA profiling of wildtype and mutant intestinal progenitor cells from intact flies exposed to normal and altered environments, as well as the identification of RNA-protein interactions critical for stem cell function.

**Summary Statement:** We report a dissection-free method to identify proximity-based RNA-protein interactions in an *in vivo* stem cell population, enabling molecular analysis of these cells at unprecedented speed and resolution.

## Introduction

The adult *Drosophila* intestine has become a premier model for understanding the biology and behavior of resident stem cells in their native context (Li and Jasper, 2016). One key approach has been transcript profiling that characterizes the gene expression signatures of cell types in both homeostatic and perturbed conditions (see, for example, (Dutta et al., 2015; Hung et al., 2020)). This approach has relied on the manual dissection of intestines. Manual dissection is not only laborious and time-consuming, but also precludes certain types of analysis that require large amounts of intact, non-degraded starting material, such as immunoprecipitation (IP). Methods that rely upon the sequencing of immunoprecipitates associated with RNA-binding proteins (RBPs) to analyze post-transcriptional control mechanisms (e.g. RiboTag) are not currently feasible in adult Drosophila intestinal stem cells but have been critical for characterizing identity and differentiation mechanisms in other stem cell lineages (Baser et al., 2019; Sanz et al., 2009; Tahmasebi et al., 2018). There is therefore a need for improved methods to profile this stem cell population in Drosophila.

Manual dissection of intestines has been required because of the lack of tools to exclusively label and manipulate individual intestinal cell types. Such tools would enable the use of the whole fly as starting material, allowing for the rapid production of large amounts of starting material. GAL4 drivers are available that label the various individual cell types in the intestinal stem cell lineage, including the intestinal stem cells (ISCs) themselves as well as their transient progenitor cell daughters, termed enteroblasts (EB), and two differentiated cell types, enterocytes (ECs) and enteroendocrine cells (EEs) (Jiang and Edgar, 2009; Micchelli and Perrimon, 2006; Zeng et al., 2010). Such GAL4 drivers can be used to express epitope-tagged transgenic proteins specifically in these various intestinal cell types that can be immunopurified. However, these GAL4 drivers are either known or likely to be active elsewhere in the adult (Biteau et al., 2010).

Here, we describe a method that uses a new intestinal progenitor GAL4 driver that is not expressed outside the intestine to profile the general transcriptome as well as specific RBP cargoes expressed in this cell type. We reasoned that an intersectional approach could be used to design such a GAL4 driver. Intersectional methods limit transgenic expression to cells in which two different enhancers are both active. In a recombinase-mediated intersectional method, for example, the recombinase under the control of one enhancer activates a GAL4 driver under the control of a second enhancer via recombinase-mediated removal of an intervening stop cassette (Fig 1A). Such a recombinase-based method involving a pan-intestinal enhancer and a progenitor enhancer should limit expression to only intestinal progenitor cells. We designed our transgenes with the KD recombinase (KDR), which was recently shown to mediate the excision, or “kick-out”, of sequences between KDR target (KDRT) sites in *Drosophila* cells (Nern et al., 2011). We chose to use KDR so that the resulting system could be used in tandem with other recombinases, like FLP, for additional manipulation of intestinal cells. Because this system is designed for intestine-specific activation of a “kick-out” transgene, we referred to it as Intestinal-KiCK-ouT, or I-KCKT.

**Figure 1:**
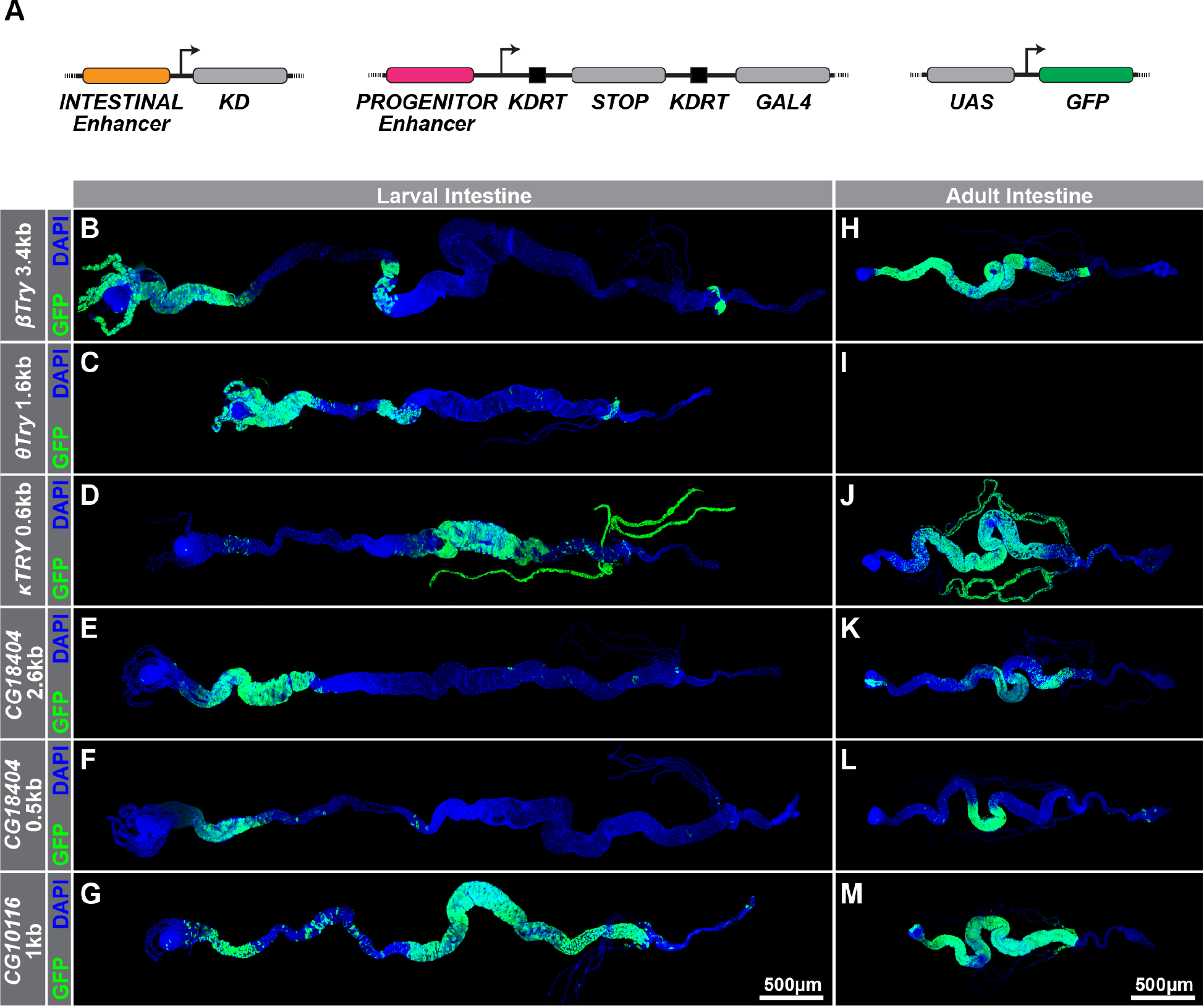
*KD* driver analysis identifies a pan-intestinal enhancer in adults. (A) Schematic of the I-KCKT system, which involves three transgenes: (i) a KD-expressing transgene under the control of an intestinal enhancer, (ii) a transgene in which a KDRT-flanked stop-cassette separates GAL4 from a progenitor enhancer, and (iii) a UAS responder, in this case, controlling GFP expression. (B-M) KD recombinase-mediated labeling of (B-G) larval and (H-M) adult intestinal cells. KD lines contain (B,H) 3.4kb *βTry*, (C, I) 1.6kb *θTry*, (D, J) 0.6kb *κTry*, (E, K) 2.6kb *CG18404*, (F, L) 0.5kb *CG18404*, or (G, M) 1kb *CG10116* enhancer sequences. KD expression pattern was detected based on KDRT-mediated activation of a *tubulin-based GAL4.p65* driver in combination with a *UAS-6XGFP* responder. Full genotypes are listed in Table S4.

## Results

### *I-KCKT-GAL4* labels most intestinal progenitor cells

To build the transgenes needed for our system, we first searched for defined pan-intestinal and progenitor enhancer sequences. Progenitor enhancer sequences have previously been identified in the *miranda* (*mira*) gene locus (Bardin et al., 2010), but the availability of a defined pan-intestinal enhancer was less clear. Transgenes with regulatory sequences from some intestine-specific genes have been reported (e.g. *mex1-GAL4, npc1b-GAL4*, etc. (Phillips and Thomas, 2006; Voght et al., 2007)), but careful inspection found that these transgenes were either not expressed throughout the intestine or in all cell types. We therefore took a candidate gene approach to identify intestine-specific enhancers based on gene expression data reported in FlyAtlas 2 (Leader et al., 2018), testing DNA fragments from intestine-specific genes for intestinal activity. KDR transgenes containing putative enhancer sequences from *βTrypsin* (*βTry*), *θTrypsin* (*θTry*), *κTrypsin* (*κTry*), *CG10116*, or *CG18404* were tested for intestinal activity in flies also harboring two additional transgenes, a *tubulin-KDRT-stop-KDRT-GAL4.p65* “kickout” transgene and a *UAS-6XGFP* responder (Shearin et al., 2014). GFP expression was monitored in both the larval and adult intestine for all resulting strains except for *θTry;* in its case, only larvae were analyzed because adults of the proper genotype failed to eclose. While all strains displayed some GFP expression in the larval intestine, most GFP patterns were patchy and non-uniform (Fig 1B-G). In the adult intestine, however, two KDR lines, *βTry-KDR* and *CG10116-KDR*, drove expression throughout the tissue (Fig 1H, M). Careful inspection of these two indicated that *CG10116-KDR* was expressed in most cells throughout the midgut, raising the possibility that the associated enhancer fragment was pan-intestinal-specific and could be used to exclusively label adult intestinal progenitor cells.

To test this possibility, we evaluated *CG10116-KDR* activity in the intestinal progenitor cells of flies harboring a *mira*-containing stop cassette transgene as well as *UAS-stinger-GFP*, a nuclear-localized GFP reporter (Barolo et al., 2000). For these experiments, we analyzed strains containing two different *mira* transgenes, one encoding an enhanced version of GAL4 with the p65 transcriptional activation domain (*mira-KDRT-stop-KDRT-GAL4.p65*) and one with a non-modified version of GAL4 (*mira-KDRT-stop-KDRT-GAL4*) that could be used in conjunction with the GAL80-dependent temporal and regional gene expression targeting (TARGET) system for conditional expression (McGuire et al., 2004). For brevity, we refer to the strains combining *CG10116-KDR* with *mira-KDRT-stop-KDRT-GAL4.p65* or *mira-KDRT-stop-KDRT-GAL4* as *I-KCKT-GAL4.p65* or *I-KCKT-GAL4*, respectively. To analyze *CG10116-KDR*-mediated expression in these strains, we compared GFP expression with that of a previously generated and validated progenitor reporter, *mira-His2A.mCherry.HA* (Miller et al., 2020), in each of the five intestinal regions (Fig 2A-J). Quantification of the percentage of mCherry+ cells that were also GFP+ indicated that ~100% of progenitor cells were labeled in most regions of both *I-KCKT* strains (Fig 2P, Q). The only two exceptions to this trend were regions 1 and 3 of *I-KCKT-GAL4.p65* intestines, where only 58.7±15.5% (n=5) and 64.4±17.0% (n=5) of progenitor cells were labeled, respectively. Importantly, we also found that no GFP+ cells were mCherry-in either strain, indicating that no non-progenitor cells were labeled in *I-KCKT* strains. To confirm this analysis, we also compared GFP expression driven by *I-KCKT-GAL4* to a second, validated progenitor reporter, *esg-LacZ* (Micchelli and Perrimon, 2006), and found similarly that almost all LacZ+ cells were also GFP+ and that no GFP+ cells were LacZ-(Fig S1). The only exception was region 3, where ~69.9±9.0% of LacZ+ cells were GFP+. This discrepancy may reflect that *esg-LacZ* labels a small subset of EEs in this midgut area (Hung et al., 2020) Collectively, this analysis indicated that almost all progenitor cells, but few if any other cells, were labeled in the intestines of *I-KCKT* strains.

**Figure 2:**
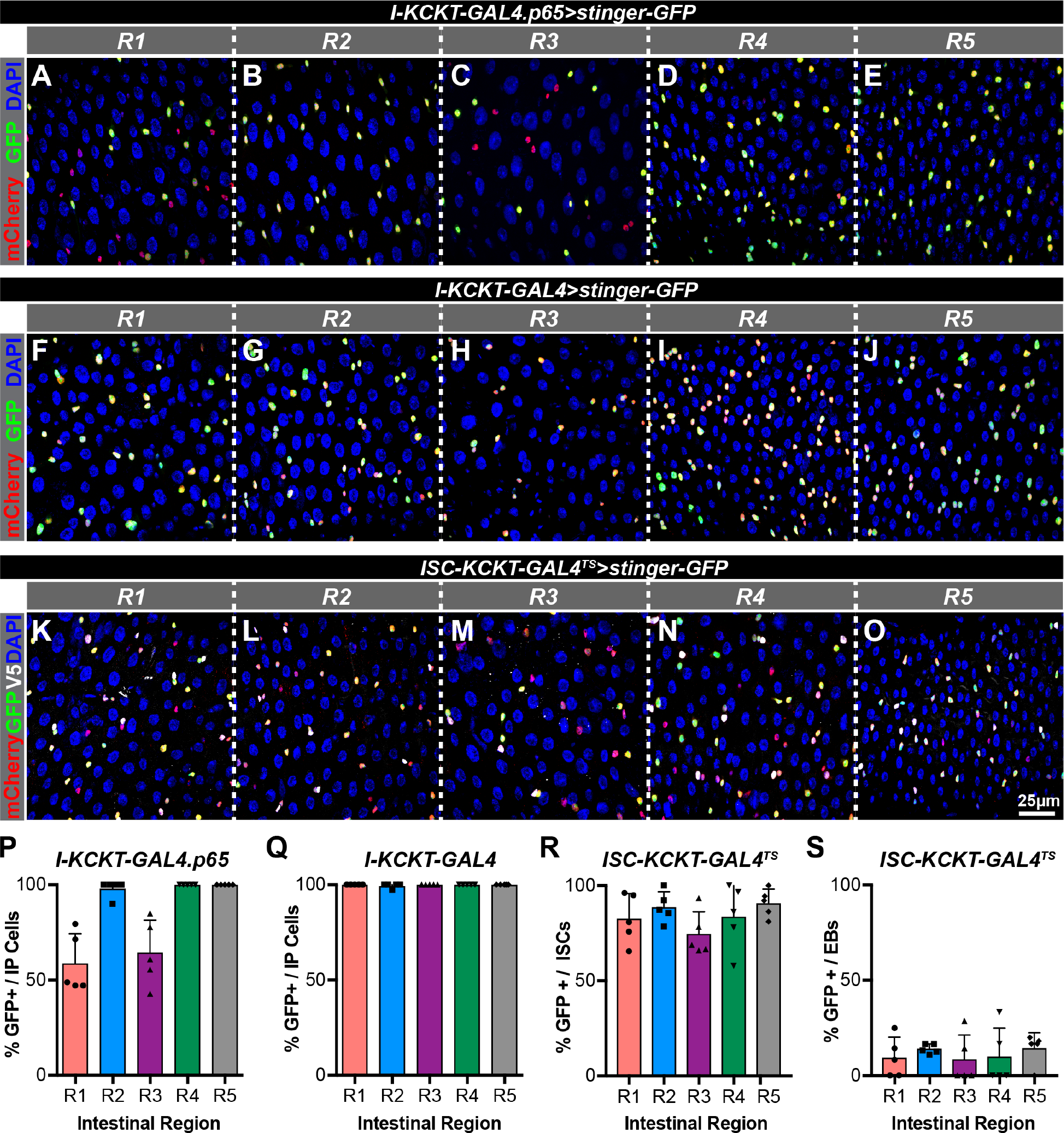
*I-KCKT-GAL4.p65* and *I-KCKT-GAL4* label most intestinal progenitor cells and *ISC-KCKT-GAL4^TS^* labels most ISCs. (A-O) Micrographs of five intestinal regions (R1-R5) stained for *stinger-GFP* in green, the intestinal progenitor marker *mira-His2A.mCherry.HA* in red, and the DAPI DNA marker in blue. GFP expression is driven by either (A-E) *I-KCKT-GAL4.p65*, (F-J) *I-KCKT-GAL4*, or (K-O) *ISC-KCKT-GAL4^TS^*. (P-Q) Histograms showing percentage of *mira-His2A.mCherry.HA-labeled* intestinal progenitor (IP) cells per intestinal region (R1-R5) that are labeled with *stinger-GFP* driven by either (A-E) *I-KCKT-GAL4.p65* or (F-J) *I-KCKT-GAL4*. (R-S) Histograms showing the percentage of (R) ISCs and (S) EBs labeled with *stinger-GFP* driven by (KO) *ISC-KCKT-GAL4^TS^*. Full genotypes are listed in Table S4.

### *I-KCKT-GAL4* is not detected in non-intestinal tissue

To determine whether intestinal progenitor cells were exclusively labeled in *I-KCKT* adults, we performed two analyses. First, we crossed *I-KCKT* strains to *UAS-6XGFP* and visually inspected adults for GFP expression. *I-KCKT* strains displayed prominent 6XGFP fluorescence in the intestine but little, if any, elsewhere (Fig 3A, B). For comparison, we similarly analyzed three other widely used drivers known to be expressed in some or all intestinal progenitor cells: a *GawB* P-element insertion in the *escargot* (*esg*) locus (*esg-GAL4*), a *GawB* P-element insertion in the *Delta* (*Dl*) locus (*Dl-GAL4*), and a *Notch*-responsive GAL4 reporter that contains binding sites for the Grainyhead and Suppressor of Hairless transcription factors (*gbe-GAL4*) (Micchelli and Perrimon, 2006; Zeng et al., 2010). All three were detected in the intestine (Fig 3E-G); *Dl-GAL4* and *gbe-GAL4* also displayed prominent non-intestinal expression while *esg-GAL4* appeared more similar to the *I-KCKT* with regard to intestinal specificity. For a more rigorous analysis, we also performed comparative Western Blot analysis for 6XGFP expression on three different protein extracts generated from each strain: whole animal extract (total), extract from dissected gastrointestinal tracts that included the malphigian tubules (intestine), and extract from all remaining non-intestinal tissue (carcass) (Fig 3H). For both *I-KCKT* strains, GFP was detected in intestinal but not carcass extract, and the amount in the intestine was roughly similar to the amount in the total extract. In contrast, GFP was detected in both the intestine and carcass of the three comparison strains, *esg-GAL4, Dl-GAL4*, and *gbe-GAL4*. We also tested the conditional expression of *mira-KDRT-stop-KDRT-GAL4* by generating an *I-KCKT-GAL4^TS^* strain that harbored an ubiquitously expressed, temperature sensitive GAL80 transgene (*tub-GAL80^TS^*) and then performing both analyses on flies that had been incubated at the non-permissive (18°C) and permissive (30°C) temperatures (Fig 3H-J). This analysis detected no GFP expression at the non-permissive temperature, and clear expression at the permissive temperature. We noted that *I-KCKT-GAL4^TS^-driven* GFP signal can also be detected in the progenitor cells of the malphigian tubules (arrowheads in Fig 3J), which are *esg^+^* cells that are related to midgut progenitor cells (Singh et al., 2007). We also note that, like *esg-GAL4* but to a lesser degree, *I-KCKT-GAL4^TS^* lost its progenitor-specificity in the intestinal tissue of aged animals (Fig S2). Collectively, these results indicated that, unlike other commonly used progenitor drivers, *I-KCKT*-GAL4-based expression could not be detected outside the intestine.

**Figure 3:**
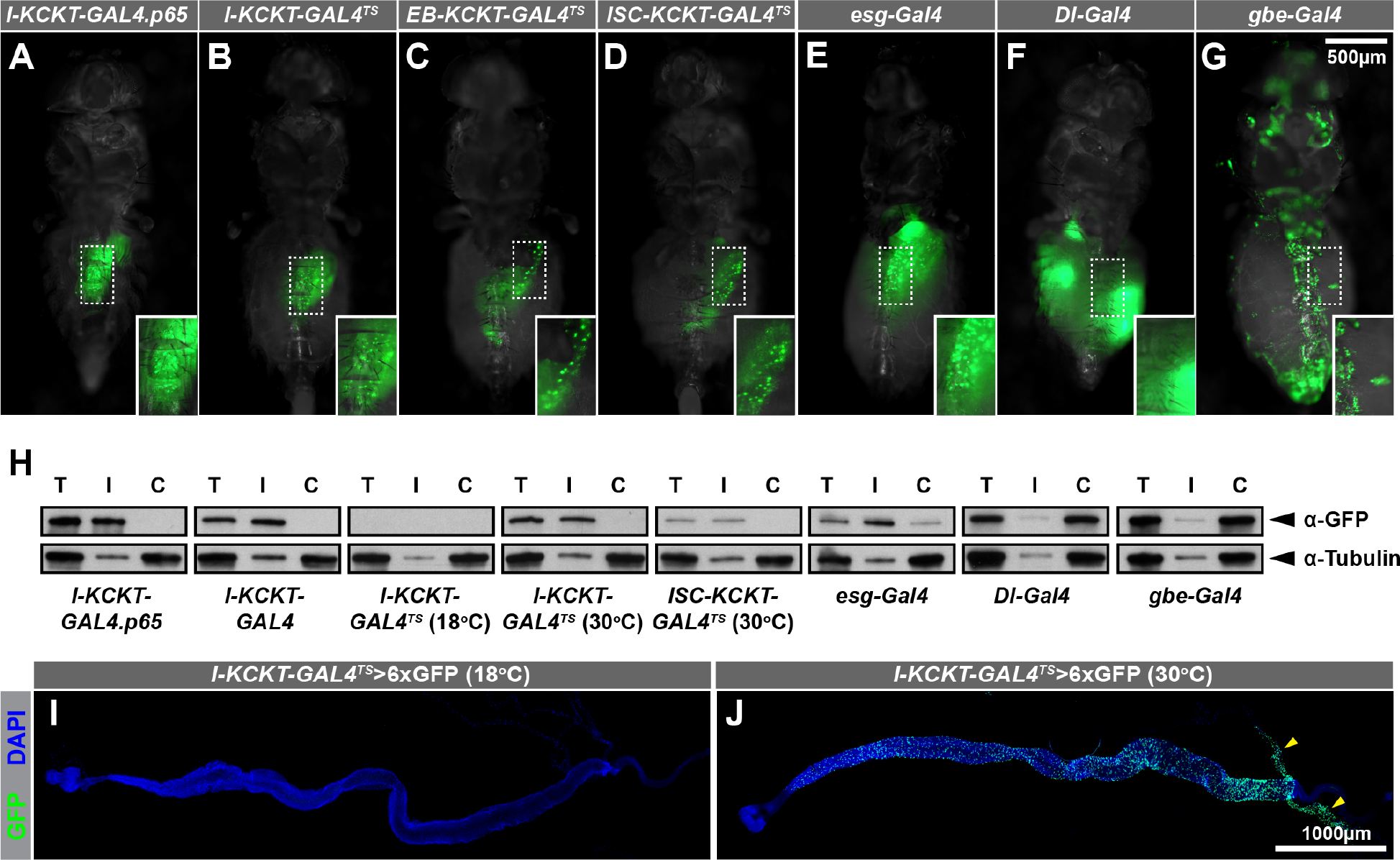
*I-KCKT-GAL4* drivers are intestine-specific. (A-G) Pictures of adult females showing *6XGFP* expression patterns driven by (A) *I-KCKT-GAL4.p65*, (B) *I-KCKT-GAL4^TS^*, (C) *EB-KCKT-GAL4^TS^*, (D) *ISC-KCKT-GAL4^TS^*, (E) *esg^P{GawB}NP5130^-GAL4*, (F) *Dl^05151-G^-GAL4*, or (G) *Su(H)GBE-GAL4*. Insets are enlargements of abdominal regions displaying intestinal GFP expression. (H) Western blot analysis of tissue extracts from total animals (T), dissected intestines (I), or dissected carcasses (C) probed with either anti-GFP (top) or anti-Tubulin (bottom) antibodies. Lysates were generated from adults harboring *UAS-6XGFP* driven by *I-KCKT-GAL4.p65, I-KCKT-GAL4, I-KCKT-GAL4^TS^* at 18°C, *I-KCKT-GAL4^TS^* at 30°C, *ISC-KCKT-GAL4^TS^* at 30°C, *esg^P{GawB}NP5130^-GAL4, Dl^05151-G^-GAL4*, or *Su(H)GBE-GAL4*. (I, J) Micrographs of intestines showing *6XGFP* expression patterns driven by *I-KCKT-GAL4^TS^* from adults kept either at (I) 18°C or (J) 30°C and counterstained for the DAPI DNA marker in blue. Yellow arrowheads indicate GFP expression at the base of the malphigian tubules. Full genotypes are listed in Table S4.

### *ISC-KCKT-GAL4* specifically labels adult intestinal stem cells

To further expand on the utility of the I-KCKT system, we investigated whether *I-KCKT*-based GAL4 expression could be limited to either the ISC or EB subsets of progenitor cells. To do so, we prepared strains in which the three *I-KCKT^TS^* transgenes were combined with a GAL4-silencing GAL80 transgene that contained either the EB-specific *gbe* synthetic enhancer (*gbe-GAL80*) or a fragment from the *Delta* locus reported to be active in ISCs (*GMR24H06-GAL80*) (Furriols and Bray, 2001; Guo et al., 2013). We referred to these strains as *ISC-KCKT-GAL4^TS^* and *EB-KCKT-GAL4^TS^*, respectively. Like *I-KCKT-GAL4^TS^*, both *ISC-KCKT-GAL4^TS^* and *EB-KCKT-GAL4^TS^* activity were intestine-specific based on visual and/or Western blot analysis (Fig 3C, D, H). To evaluate the cell type specificity of this activity, we drove GFP expression in both strains and compared it to a dual reporter combination that effectively distinguishes ISCs and EBs (Fig S3). These dual reporters are the progenitor marker *mira-His2A.mCherry.HA* (Miller et al., 2020) and the EB-specific marker *3Xgbe-smGFP.V5.nls* (Buddika et al., 2020a). For this analysis, we scored mCherry+, V5- cells as ISCs and mCherry+, V5+ cells as EBs. Most ISCs were specifically labeled in the *ISC-KCKT-GAL4^TS^* strain: 84.1±12.3% (n=5) of ISCs were labeled whereas 11.2±10.0% (n=5) of EBs were labeled across all five regions of the intestine (Fig 2K-O, R, S). In contrast, EB-specific labeling in *EB-KCKT-GAL4^TS^* was less effective: only 30.7±20.1% (n=5) of EBs were labeled whereas 57±19.8% (n=5) of ISCs were labeled across all five regions of the intestine (Fig S4). Various possibilities could explain this *EB-KCKT-GAL4^TS^* result, including that the GMR24H06 enhancer fragment was not active in most ISCs and/or that GAL80 activity perdured in EB cells. Nevertheless, this analysis indicated that *ISC-KCKT-GAL4^TS^* specifically labeled most ISCs.

### CLIP-seq of PABP using *I-KCKT-GAL4* identifies progenitor expressed genes

We next investigated whether these *I-KCKT* strains could be used to streamline current methods to molecularly profile progenitor cells. RNA-seq profiling methods that rely on *esg-GAL4*, for example, require labor-intensive dissection of hundreds of intestines followed by digestion of this tissue to release labeled progenitor cells for fluorescent activated cell sorting (FACS) (Dutta et al., 2015; Fast et al., 2020; Korzelius et al., 2019; Li et al., 2018) (Fig 4A, left). Because *I-KCKT-GAL4* activity was limited to intestinal progenitor cells, we tested whether a cross-linking immunoprecipitation and sequencing (CLIP-seq) analysis of *I-KCKT-GAL4-*driven, FLAG-tagged poly-A Binding Protein (PABP) that used whole animal lysate as its starting material would recover progenitor cell RNA (Fig 4A, right). PABP is a general mRNA-binding protein that has been used to profile the mRNA transcriptomes of other cell types (Hwang et al., 2016; Yang et al., 2005). We first verified that the FLAG-tagged version of PABP that had previously been used to profile mRNAs in *Drosophila* photoreceptor cells was expressed in progenitor cells when crossed to *I-KCKT-GAL4^TS^* (Fig S5) (Yang et al., 2005). Then, we crushed frozen flies from this strain to generate a fine powder, subjected this powder to UV crosslinking to covalently link protein and RNA complexes, immunoprecipitated PABP using anti-FLAG beads, and recovered the associated RNAs. Independent libraries were prepared from duplicate PABP immunoprecipitates (PABP IPs). In parallel, two libraries were also prepared from RNA extracted from the starting whole animal lysate, and we used these “input” libraries as our normalization controls. Differential gene expression analysis found 1,661 transcripts enriched in the PABP IP samples and 3,293 enriched in input (Fig. 4B, Table S1). Representative examples of these respective classes included the progenitor-enriched *esg* transcript and the ovary-enriched *oskar* transcript (*osk*) (Fig. 4C-D).

**Figure 4:**
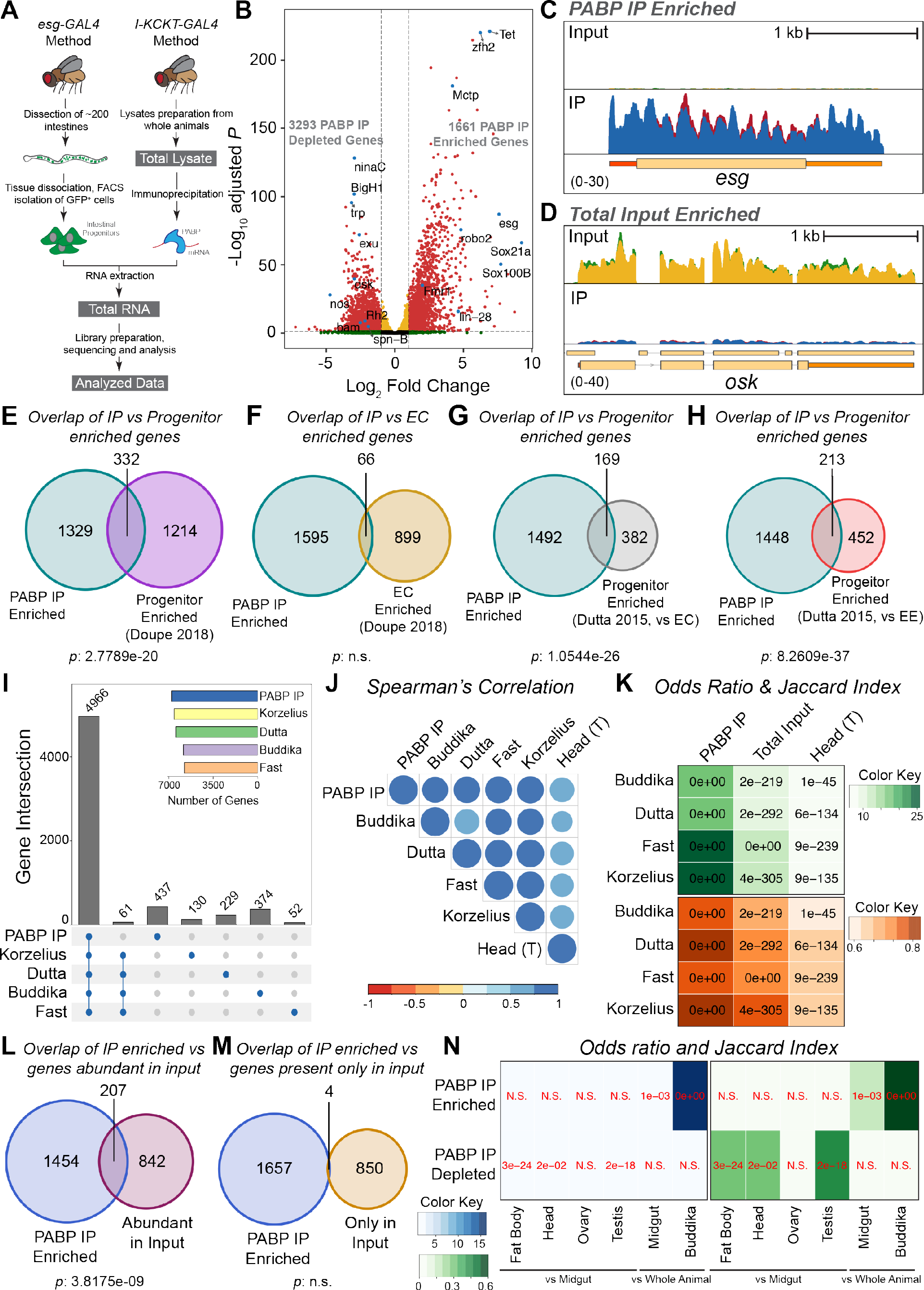
CLIP-seq of PABP driven by *I-KCKT-GAL4* identifies progenitor expressed genes. (A) Schematic representation of conventional FACS (left) and *I-KCKT*-based (right) intestinal progenitor RNA-sequencing. (B) Volcano plot of differentially expressed genes in PABP IP versus total input. Each dot represents a single gene. Yellow indicates a false discovery rate adjusted p-value (FDR) < 0.05 and a Log_2_ fold change < 1 or > −1. Green indicates Log2 fold change > 1 or < −1 and an FDR >= 0.05. Red indicates an FDR < 0.05 and log2 fold change > 1 or < −1. A selected set of significantly changed genes have been labeled in blue. (C-D) Genome browser tracks of normalized total input and PABP IP at the (C) *esg* or (D) *osk* loci. Note that the two replicates of the total input (green and yellow) and the two replicates of the PABP IP (red and blue) have been overlayed. (E-H) Venn diagrams showing genes significantly enriched in PABP-CLIP that overlap with Doupe’s progenitor enriched (E) genes, (F) Doupe’s EC enriched genes, (G) Dutta’s progenitor enriched genes as compared to EC genes, or (H) Dutta’s progenitor enriched genes as compared to EE genes. (I) Upset plot showing the overlap of moderately expressed genes identified by PABP CLIP-seq versus RNA-seq of FACS-isolated progenitor cells reported in Korzelius, Dutta, Buddika and Fast. Numbers above each bar shows the size of each intersection. (J) Correlogram of Spearman’s *rho* values for pairwise comparisons of progenitor expressed genes in PABP CLIP-seq, progenitor RNA-seq reported in Buddika, Dutta, Fast, or Korzelius, or RNA-seq from *Drosophila* female head. (K) Heatmaps of odds ratio (in green) and Jaccard index (in orange) values for pairwise comparisons of PABP IP CLIP-seq, RNA-seq of total input, or RNA-seq of dissected female heads against FACS-based progenitor RNA-seq. Numbers on each colored box show the p-value for each overlap based on Fisher’s exact test. (L-M) Venn diagrams showing genes significantly enriched in PABP-CLIP that overlap with (L) the top 10% of input genes based on normalized expression values, or (M) genes present in input but not in any of the FACS-based RNA-seq gene lists. (N) Heatmaps of odds ratio (in blue) and Jaccard index (in green) values of pairwise comparisons of PABP IP enriched or PABP IP depleted genes to six other gene sets. These sets include fat body, head, ovary or testis enriched genes (identified relative to midgut genes) as well as midgut or progenitor enriched genes from Buddika (identified relative to whole animal input). Numbers on each colored box show the p-value for each overlap based on Fisher’s exact test.

To evaluate whether the PABP IP-enriched transcripts included transcripts known to be enriched in progenitor cells, it was cross-referenced to two published datasets. First, we compared our results to transcriptome profiles obtained by Doupé, who used a DamID based method to identify progenitor-enriched and EC-enriched transcripts (Doupe et al., 2018). Twenty percent of the 1,661 were present in Doupé’s progenitor-enriched transcripts, whereas only 4% were present in the EC-enriched set (Fig. 4E-F). In addition, we re-analyzed published RNA-seq datasets from FACS-isolated progenitors, ECs, and EEs (Dutta et al., 2015), identified transcripts enriched in progenitor cells when compared to either ECs or EEs and found that 31%-32% of these enriched transcripts overlapped with transcripts enriched in the PABP IP (Fig. 4G-H, Table S2). To benchmark these comparisons with the Doupé and Dutta datasets, we performed analogous comparisons between them and an RNA-seq dataset of *esg+* progenitor cells that we generated using FACS and the same library preparation kit as for the PABP IP libraries (Buddika et al., 2020b), and found similar levels of overlap (Fig. S6A-D). This analysis indicated, therefore, that the transcripts most highly represented in the PABP IP relative to whole animal lysate included a substantial number of genes previously shown to be enriched in progenitor cells.

We next focused on comparing the entire PABP-associated transcriptome to the RNA-seq profile of FAC sorted progenitor cells, using four previously reported RNA-seq datasets. These datasets were reported in (Buddika et al., 2020b; Dutta et al., 2015; Fast et al., 2020; Korzelius et al., 2019), and we refer to them as Buddika, Datta, Fast, and Korzelius, respectively. Read counts for the four FACS datasets as well as the PABP IP dataset were normalized in parallel to facilitate direct comparison; a transcript was considered to be expressed in a dataset if it had a normalized expression value greater than 10 in each replicate. Only 61 genes found in all four FACS datasets were not present in the PABP IP dataset, indicating that the PABP IP identified 98.8% (4,966 out of 5,027) of core progenitor transcripts common to all four distinct FACS datasets (Fig 4I). An additional 1,839 transcripts were found in the PABP IP, and 76.2% of these were also found in at least one other FACS dataset. Four hundred and thirty-seven transcripts (6.4%) were unique to the PABP IP, a number in line with the numbers unique to each FACS dataset and that likely reflected differences caused by the technical approach. These could include normal deviations associated with distinct experimental settings, as well as differences related to the preparation of flash frozen versus mechanically disrupted samples. Altogether, these results indicated that the PABP IP method effectively identified progenitor transcripts.

To evaluate the significance of the overlap between these five datasets, Spearman correlation matrices were generated and visualized as correlograms. As a negative control for this analysis, we also prepared a sixth dataset of head tissue data from FlyAtlas 2 (Leader et al., 2018). As expected, the PABP IP correlated well with all FACS based progenitor datasets while, furthermore, all five of these datasets showed a similar, lower correlation with the non-intestinal dataset (Fig. 4J). We then used the GeneOverlap R package to perform Fisher’s exact test to evaluate the statistical significance of the overlap between different datasets. The Fisher’s exact test also computes both an odds ratio and a Jaccard index, which represents the strength of association and similarity between two datasets, respectively. The PABP IP scored highly in both these indices when compared to each of the four FAC-sorted datasets and significantly higher than either the total input or head tissue datasets, further indicating the similarity between the PABP IP and FACS-based datasets (Fig. 4K).

Finally, we evaluated whether the PABP IP contained progenitor-enriched transcripts that we identified from systematic re-analysis of the four FACS-based datasets. For this re-analysis, differential expression analysis was used to select for transcripts enriched in Buddika, Datta, Fast, and Korzelius compared to two sources of whole animal data, total input from the PABP experiment or reanalyzed whole female adult dataset from FlyAtlas 2. The PABP IP was then compared to each of these computed progenitor enriched datasets. As expected, PABP IP enriched genes showed significant low p-values and high odds ratio and Jaccard index scores, strengthening the positive relationship of PABP and FACS-based datasets (Fig. S6E). Altogether, this bioinformatic analysis supported the conclusion that the PABP IP dataset represented the gene expression profiles of progenitor cells.

### CLIP-seq of I-KCKT-GAL4-driven PABP has little background

Having established that the PABP IP dataset contained progenitor genes, we also sought to evaluate how much non-specific background was represented in this dataset. Such noise might come from the post-lysis association of transgenic PABP with mRNAs from non-progenitor cells (Mili and Steitz, 2004). Hypothesizing that the most abundant transcripts in the whole animal lysate might be most prone to associate with PABP post-lysis, we first identified the top 10% of genes in the input based on the normalized expression values and found 1049 genes. Only 207 of these genes were also identified in the PABP IP enriched gene set, suggesting that transcripts abundantly expressed in the starting lysate were not non-specifically recovered in the IP (Fig. 4L). Supporting this observation, we identified 854 genes in the input that were expected not to be present in the PABP IP (Fig. S6F) and confirmed that only 4 of these transcripts were present in the PABP IP enriched gene list (Fig. 4M). Finally, we again used reanalyzed FlyAtlas 2 data to identify genes enriched in fat body, head, ovary and testis relative to midgut, and then compared those tissue-enriched genes to the genes either enriched in or depleted from the PABP IP. We found that odds ratio and Jaccard index scores were low for PABP-enriched genes, indicating minimal overlap, and significantly higher for PABP-depleted genes (Fig. 4N). As a positive control for this analysis, we performed the same comparisons with genes enriched in the Fly Atlas 2 midgut dataset as well as genes enriched in the Buddika FACS dataset. As expected, PABP IP enriched genes show highest association with genes enriched in FACS-isolated progenitors, and significant overlap was also observed with the midgut enriched gene list (Fig. 4N). Collectively, this analysis indicated that the PABP dataset contained minimal background noise from other tissues, likely due to the stringency of the washes after the crosslinking step.

### *I-KCKT*-based eCLIP analysis identifies FMRP-bound mRNAs in the intestine

To illustrate the breadth of its possible applications, we employed *I-KCKT-GAL4* to identify the RNA cargo of a more selective RBP, Fragile X Mental Retardation Protein (FMRP), specifically in progenitor cells using enhanced Crosslinking Immunoprecipitation sequencing (eCLIP-seq). FMRP limits ISC expansion but its mRNA targets in these cells are unknown (Luhur et al., 2017). The eCLIP method contains a number of key modifications in comparison to other CLIP methods, including adapter ligation steps that map the exact position of crosslinking and the preparation of a size-matched input (SMI) control sample for stringent normalization (Van Nostrand et al., 2016). We first verified that *UAS-FMRP.FLAG.GFP* in combination with *I-KCKT-GAL4^TS^* was detected in progenitor cells after two and ten days of induction and that it caused a reduction in progenitor cell number when induced for ten but not two days (Fig S5). We then prepared FMRP IP and SMI libraries from whole animals collected at the two-day timepoint (Fig S7A-B). After removing rRNA reads, multimapping reads, and duplicated reads, about 8% (8,438,543), 5% (2,487,118) and 7% (2,545,165) of total reads were recovered as usable reads from the SMI and two IP samples, UV1 and UV2, respectively (Fig S7C), consistent with the recovery rate from other eCLIP analyses (Van Nostrand et al., 2016).

Peak calling analysis identified 13,297 reproducible FMRP binding sites across 1,829 genes (Table S3) and indicated that there was high correlation between replicates (Fig S7D), demonstrating the robustness of our modified eCLIP method. Analysis of peak distribution indicated that >85% of FMRP binding sites were in protein-coding transcripts and, within these transcripts, the majority of sites were in coding sequences (Fig 5A, B). Both of these results were consistent with previous characterizations of FMRP distribution in multiple species, including from proximity-based analyses in mice and human tissues as well as activity-based analysis in fly cells (Darnell et al., 2011; Li et al., 2020; McMahon et al., 2016). In addition, we noted that this analysis identified several confirmed direct targets of Drosophila FMRP found by genetic or targeted biochemical means, including *CaMKII, chic, Dscam1, futsch, ninaE, rg* and *tral* (Cvetkovska et al., 2013; Kim et al., 2013; Monzo et al., 2010; Reeve et al., 2005; Sears et al., 2019; Sudhakaran et al., 2014; Wang et al., 2017; Zhang et al., 2001). As the first CLIP-based assay to characterize Drosophila FMRP, this analysis also identified CAUUG(A/U) as its top binding motif (Fig 5C), consistent with prior identification of the AUUG sequence in one of the top binding motifs of human FXR1 (Feng et al., 2019).

**Figure 5:**
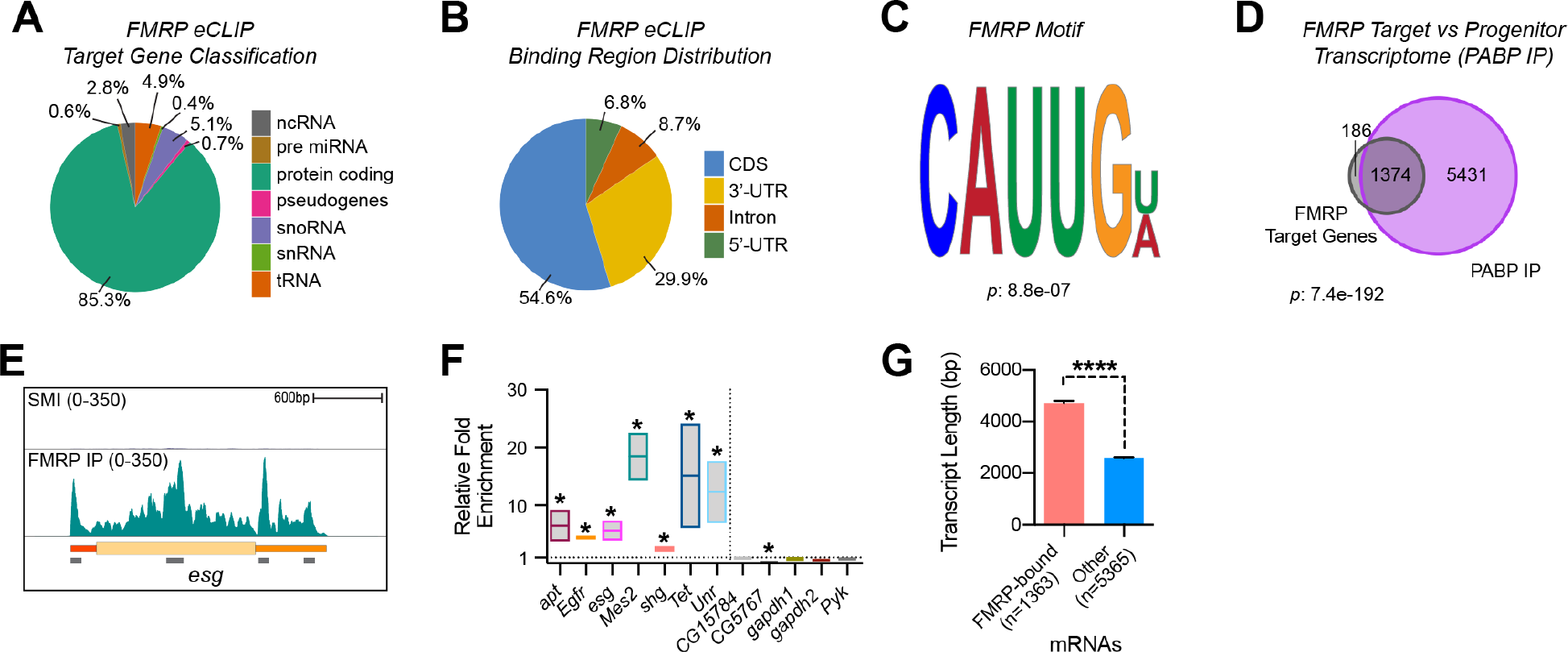
eCLIP of FMRP driven by *I-KCKT-GAL4* identifies intestinal target mRNAs. (A-B) Pie chart representing the percentage of (A) target gene types and (B) binding site distribution in mRNAs identified in FMRP eCLIP. (C) The top FMRP binding motif identified using DREME. (D) Venn diagram showing overlap between FMRP target genes with the progenitor transcriptome (identified by PABP IP). (E) Genome browser tracks of normalized SMI and FMRP IP at the *esg* locus of the *Drosophila* genome. Note that the two replicates of FMRP IP were merged prior to genome browser visualization. Locations of four FMRP binding regions are shown with grey bars. (F) Bar plot showing fold enrichment of twelve mRNAs in FMRP IP compared to whole intestinal input as determined by qPCR. Negative controls are separated from the other genes by a dotted line. (G) Bar plot showing the mean transcript length of FMRP-bound versus non-bound transcripts that are expressed in intestinal progenitors. Note that the longest transcript isoform was used for this analysis whenever a gene had multiple transcript isoforms. *p < 0.05; ****p < 0.0001.

The FMRP mRNA cargo includes ~20% of the protein-coding transcripts expressed in progenitor cells that were identified by the PABP IP analysis described above (Fig 5D). These FMRP targets were significantly associated with processes related to stem cell proliferation, stem cell maintenance, cell differentiation, and translation repressor activity (Fig. S7E), consistent with the known roles of FMRP in stem cell populations such as ISCs (Luhur et al., 2017). One representative example of these FMRP targets was *esg* (Fig 5E), which plays critical roles in progenitor cells where it is a known target of post-transcriptional control (Antonello et al., 2015; Korzelius et al., 2014). We also noted that ~12% of the FMRP cargo are from genes not identified in either PABP IP or progenitor cell RNA-seq (Fig 5D, S7F). This signal could therefore represent post-lysis association of FMRP.FLAG.GFP with non-progenitor mRNAs. However, none of 186 FMRP target genes that were not represented in the PABP IP were among the 1049 most abundant transcripts present in the whole fly input (Fig S7G). We therefore favor the hypothesis that these FMRP targets represent weakly-expressed progenitor transcripts that were not recovered in the PABP IP because of their low level of expression.

To verify these eCLIP-identified FMRP targets, we first performed qPCR on RNA immunoprecipitated with endogenous FMRP from wildtype intestines (Fig S7H). We chose seven target genes (*apt*, *Egfr, esg, Mes2, shg, Tet*, and *Unr*) as well as five negative controls (*CG15784, CG5767, gapdh1, gapdh2*, and *Pyk*) for this analysis. As expected, the FMRP target genes showed between 2- and 18-fold enrichment in FMRP IP relative to whole intestine input, while the negative controls were either not significantly enriched or significantly de-enriched in the FMRP IP (Fig 5F). In addition, we found that the length distribution of the FMRP-bound mRNAs was significantly longer than the other protein-coding transcripts expressed in progenitor cells (Fig 5G), consistent with recent ribosome foot printing that showed that FMRP preferentially regulates the translation of large proteins in Drosophila oocytes (Greenblatt and Spradling, 2018). Altogether, these results indicated that applying eCLIP to whole animal tissue that contained progenitor cells that expressed FLAG-tagged FMRP effectively recovered the FMRP cargo from these cells.

## Discussion

Here we describe a method to purify the RNA cargoes associated with intestinal progenitor RBPs using whole animal lysate as the starting material. Because it is GAL4-based, this method will have wide applicability for profiling the general PABP-bound transcriptome of wildtype and, when coupled with UAS-RNAi transgenes, mutant progenitor cells. In addition, it opens the possibility of molecularly characterizing the stem cell targets of selective RBPs, a growing number of which have been shown to play critical roles in progenitor cells (e.g. LIN28, FMRP, SPEN, TIS11 (Andriatsilavo et al., 2018; Chen et al., 2015; Luhur et al., 2017; McClelland et al., 2017)). Finally, this method can be used in conjunction with additional GAL4-based mRNA profiling methods, such as TU-tagging and ribosome profiling (Chen and Dickman, 2017; Hida et al., 2017), as well as methods that target other classes of RNAs, including microRNAs (Luhur et al., 2013). While the method is currently limited to progenitor cells, its applicability could be further expanded by generating kickout transgenes that drive expression in additional intestinal cell types such as EEs and ECs. Furthermore, while we focus on protein-RNA interactions in this study, we also expect that *I-KCKT-GAL4* in combination with mass-spectrometry can be used to identify protein-protein interactions and thereby characterize protein complexes in progenitor cells.

The I-KCKT method offers a number of advantages over current progenitor cell profiling methods, which typically involve the FAC sorting of progenitor cells from dissected and then mechanically disrupted intestines. The use of whole animals rather than dissected intestines as starting material greatly expedites sample preparation, enabling the analysis of larger numbers of samples that could be used to test additional conditions and manipulations. In addition, because sample preparation is fast, this approach may effectively capture labile RNA profiles. Along these lines, the elimination of the FAC sorting step will also likely improve the accuracy of results, since FAC sorting is known to affect gene expression profiles likely due to the time and mechanical disruption associated with this step (Andra et al., 2020; Richardson et al., 2015). In addition, this method could make molecular profiling of progenitors more accessible, since cell sorting runs can be costly and the equipment is not always available.

While the conditional expression of transgenic PABP to retrieve bulk mRNA has been used to approximate the transcriptomes of specific cell types in a variety of species (Roy et al., 2002; Tenenbaum et al., 2000; Yang et al., 2005), some issues regarding PABP should be considered when using this approach. PABP displays some differential binding affinity to poly(A) RNAs that can introduce bias (Yang et al., 2005). In addition, retrieval of RNAs with poly(A) tails misses deadenylated mRNAs as well as some noncoding RNAs, thereby biasing resulting datasets towards actively translating coding transcripts. Finally, the cell toxicity associated with transgenic expression of PABP likely alters endogenous gene expression. Nevertheless, the cargo of PABP is better reflective of cell transcriptomes than the cargos of other general RNA binders and, furthermore, can be analyzed with paperCLIP to finely map 3’UTR ends and uncover alternative polyadenylation sites in a cell type specific manner (Hwang et al., 2016; Tenenbaum et al., 2000). It should also be noted that PABP IP results are well suited for comparison with RNA-seq libraries that are generated via poly(A)-selection, rather than rRNA depletion, since both methods select for polyadenylated transcripts.

As the first CLIP-based analysis of Drosophila FMRP, this study reports both novel mRNA targets as well as the binding motif of this protein in intestinal progenitor cells. The set of FMRP targets identified here displays partial overlap with those identified in two recent studies that used either ribosome profiling or a proximity-based activity assay to identify FMRP targets in oocytes, cultured cells, or neurons (Greenblatt and Spradling, 2018; McMahon et al., 2016). Differences in the targets reported by these studies likely reflect not only their distinct technical approaches but also the variety of cell types analyzed, since FMRP is known to display cell type binding preferences in other species (Maurin et al., 2018). Because loss of FMRP leads to Fragile X Syndrome (FXS), the leading form of human intellectual disability in humans, analysis of FMRP has focused on its activity in differentiated neurons. However, FMRP also functions in stem cell populations, and its dysregulation in these cells likely contribute to understudied FXS symptoms including elevated brain size, accelerated growth, and gastrointestinal problems (Luhur et al., 2017). Thus, future analysis of the stem cell targets of FMRP identified here may characterize regulatory relationships that are relevant to therapies designed to treat the entire repertoire of FXS symptoms.

A current key limitation of the I-KCKT method is its purification of epitope-tagged UAS-based transgenic protein rather than endogenous protein. This raises the concern that the transgenic protein might be expressed at higher-than-endogenous levels, which might cause inappropriate interactions with non-target mRNAs. Because the TARGET system controlling transgene expression is temperature sensitive, this concern can be addressed by testing multiple temperatures in the permissive range to identify conditions supporting physiological protein expression. As an alternative approach to address this concern, our current effort is focused on modifying endogenous RBP loci to contain FRT-flanked epitope tags. I-KCKT-based expression of FLP should then lead to the production of epitope-tagged endogenous protein in progenitor cells that would be available for eCLIP-based analysis. Thus, we expect that I-KCKT-based methods will allow unprecedented analysis of RNA-based mechanisms of progenitor cells.

## Materials and Methods

### Drosophila strains and husbandry

All fly strains were cultured on standard Bloomington *Drosophila* stock center media (https://bdsc.indiana.edu/information/recipes/bloomfood.html). Flies were reared in 18°C, 25°C and 29°C incubators set for a 12hr light/dark schedule and 65% humidity. The genotypes of all strains used in this study are listed in Table S4. Transgenesis to create new strains was performed by Rainbow Transgenic Flies using plasmid DNA described below. Additional stocks were obtained from the Bloomington Drosophila Stock Center (*{20XUAS-6XGFP}attP2, P{UAS-Stinger}2, P{tubP-GAL80[ts]}20, P{UAS-dPF}D*), Steven Hou (*P{Su(H)GBE-GAL4, P{GawB}Dl^05151-G^*), the Kyoto Drosophila Stock Center (*P{GawB}NP5130*), and Steve Stowers (*{20XUAS-DSCP-6XGFP}attP2*). For GAL80-dependent conditional expression experiments, flies were reared at 18°C, collected over 2 days, and then shifted to 30°C for up to 10 days before analysis. The following strains have been deposited at the Bloomington Drosophila Stock Center: (**i**) *I-KCKT-GAL4.p65* (BDSC #. Full genotype: *mira-KDRT>-dSTOP-KDRT>-GAL4.p65}attP40*; *{CG10116-KD.PEST}attP2*), (**ii**) *I-KCKT-GAL4^TS^* (BDSC #. Full genotype: *{mira -KDRT>-dSTOP-KDRT>-GAL4}attP40, P{tubP-GAL80[ts]}20; {CG10116-KD.PEST}attP2*), (**iii**) *ISC-KCKT-GAL4^TS^* (BDSC #. Full genotype; *{gbe-GAL80}ZH-2A; {mira -KDRT>-dSTOP-KDRT>-GAL4}attP40, P{tubP-GAL80[ts]}20; {CG10116-KD.PEST}attP2*).

### Transgenes

*KD enhancer transgenes:* PCR products containing enhancer fragments amplified from genomic DNA were cloned into the HindIII and AatII sites of *pJFRC161* (*20XUAS-IVS-KD.PEST*, a gift from Gerald Rubin, Addgene plasmid 32140) using HiFi DNA Assembly Mix (New England Biolabs). Oligo pairs used to amplify enhancer fragments from *βTry* (3437/3438), *θTry* (3439/3440), *κTry* (3446/3447), *CG18404* (3443/3444 for the 2.6 kb fragment, 3445/3443 for the 0.5kb fragment), and *CG10116* (3441/3442) are shown in parentheses, and oligo sequences are reported in Table S5. Junctions of resulting plasmids were verified by sequencing prior to the preparation of plasmid DNA for transgenesis. *KD* enhancer transgenes were inserted into the *attP2* landing site.

*KDRT GAL4 transgenes: pJFRC164* (*21XUAS-KDRT>-dSTOP-KDRT>-myr.RFP*, a gift from Gerald Rubin, Addgene plasmid 32141) was used as a backbone for KDRT-flanked stop cassette plasmids. Four-way HiFi DNA Assembly reactions were performed with (i) 7.2kb HindIII/XbaI-digested *pJFRC164* plasmid backbone, (ii) 1.4kb AatII/XhoI-digested *pJFRC164* KDRT>-dSTOP-KDRT fragment, (iii) PCR-amplified GAL4-containing fragments, and (iv) PCR amplified enhancer-containing fragments. GAL4.p65 was amplified from *pBPGAL4.2.p65Uw* (a gift from Gerald Rubin, Addgene plasmid 26229) with oligo pair 3433/3434, while GAL4 sequence was amplified from *pBPGAL4.1Uw* (a gift from Gerald Rubin, Addgene plasmid # 26226) with oligo pair 4401/4402. Enhancer fragments include: (i) 2.6 kb *tubulin* fragment amplified with oligo pair 3626/3627, (ii) 2.6 kb 5’ and 1.6 kb 3’ *miranda* fragments amplified with oligo pairs 3489/3490 and 3491/3492, respectively, and (iii) 3.4 kb *delta* fragment amplified with oligo pair 3431/3432. Plasmid junctions and open reading frames were verified by sequencing prior to the preparation of DNA for transgenesis. Oligo sequences are reported in Table S5, and complete plasmid sequences are available upon request. KDRT transgenes were inserted into the *attP40* landing site.

*GAL80 transgenes:* For *GMR24H06-GAL80*, the R24H06 enhancer fragment was PCR amplified from genomic DNA and subcloned upstream of the GAL80 open reading frame in a *pATTB*-containing transformation plasmid. *3xgbe-GAL80* was generated in a similar manner except using oligos encoding three tandem copies of *gbe* sequence (Furriols and Bray, 2001). Both of these *GAL80* transgenes were inserted into the *ZH-2A* landing site.

### Dissections and immunostaining

Adult female flies were dissected in ice cold 1xPBS and fixed in 4% paraformaldehyde (Electron Microscopy Sciences) in PBS for 45 min. For all samples to be stained with antibodies other than anti-Dl, tissue was then washed in 1xPBT (1xPBS, 0.1% Triton X-100) and then blocked (1xPBT, 0.5% Bovine Serum Albumin) for at least 45 min. Samples were incubated at 4°C overnight with primary antibodies, including rabbit anti-RFP (Clontech, 1:1000), mouse anti-V5 (MCA1360GA, Bio-Rad, 1:250), chicken anti-LacZ (ab9361, Abcam, 1:2000), mouse anti-FLAG (F1804, Sigma, 1:1000), and rabbit anti-HA (3724S, Cell Signaling Technology, 1:1000). The following day, samples were washed in 1xPBT and incubated for 2-3 hours with secondary antibodies, including AlexaFluor-488 and −568-conjugated goat anti-rabbit, -mouse, -rat and - chicken antibodies (Life Technologies, 1:1000). Finally, samples were washed in 1xPBT, including one wash with DAPI (1:1000), and mounted in Vectashield mounting medium (Vector Laboratories).

For staining with anti-Dl antibodies, an alternative sample preparation scheme adapted from (Lin et al., 2008) was followed. Briefly, intestines were dissected in ice-cold Grace’s Insect Medium (Lonza Bioscience) and fixed in a 1:1 (V/V) mixture of heptane (Sigma) and 4% paraformaldehyde (Electron Microscopy Sciences) in water for 15 min. The bottom aqueous paraformaldehyde layer was removed, 500μl of ice-cold methanol was added, and shook vigorously for 30 seconds. The methanol/heptane mixture was removed and incubated with 1ml ice-cold methanol for 5 min. Next, samples were gradually rehydrated with a series of 0.3% PBT (1xPBS, 0.3% Triton X-100):Methanol (3:7, 1:1, 7:3) washes, washed with 0.3% PBT alone for another 5 min and blocked (0.3% PBT, 0.5% Bovine Serum Albumin) for at least 45 min. The primary antibody was mouse anti-Delta (C594.9B, Developmental Studies Hybridoma Bank [DSHB], 1:500), and secondary antibodies are described above. Samples were mounted in ProLong Diamond mounting medium (Invitrogen).

### Microscopy and image processing

Images of whole flies were collected on a Zeiss Axio Zoom microscope and images of dissected intestines were collected on a Leica SP8 Scanning Confocal microscope. Samples to be compared were collected under identical settings on the same day, image files were adjusted simultaneously using Adobe Photoshop CC, and figures were assembled using Adobe Illustrator CC.

### Western blot analysis

Female *Drosophila* flies were used for protein isolation. Extract was prepared from either whole animals or separately from the intestines and remaining carcass tissue of dissected animals. Tissues were lysed in protein lysis buffer (25mM Tris-HCl pH 7.5, 150mM KCl, 5mM MgCl2, 1% NP-40, 0.5mM DTT, 1x Complete protease inhibitor cocktail (Sigma)), protein extracts were resolved on a 4-20% gradient polyacrylamide gel (Bio-Rad), transferred to Immobilon-P membrane (Millipore) and probed with rabbit anti-GFP (ab290, Abcam, 1:10,000) or mouse anti-a-tubulin (12G10, Developmental Studies Hybridoma Bank, 1:1000) antibodies. For IP verification, blots were stained with mouse anti-FLAG (F1804, Sigma, 1:3500) or mouse anti-FMRP (5A11, Developmental Studies Hybridoma Bank, 1:500). Subsequently, blots were washed extensively with 1xTBST (1xTBS, 0.1% Tween-20) and incubated with anti-rabbit or -mouse conjugated HRP secondary antibodies. After extensive secondary washes with 1xTBST, blots were treated with ECL-detection reagents (Thermo Scientific) and finally exposed to chemiluminescence films (GE Healthcare).

### RNA immunoprecipitation, library construction, and sequencing

Whole flies were collected and immediately snap frozen in liquid nitrogen and stored at - 80°C until sample preparation. Two hundred flies (100 male and 100 female) of each sample were ground into a fine powder using a pre-cooled mortar and pestle on dry ice. The powder was irradiated three times at 120 mJ per cm2 in a UV cross-linker (Stratagene), with mixing between each irradiation cycle to maximize surface exposure. The fly powder was transferred to a 2ml tube containing 1ml lysis buffer (150mM NaCl, 50mM Tris-HCl, pH7.5, 1mM EDTA, 1% Triton X-100, 1% Sodium Deoxycholate, 0.1% SDS, 5X Complete Protease Inhibitor Cocktail (Sigma, 11836145001), 4U/ml RNase Inhibitor Murine (BioLabs, M0314L) using a cold spatula, and the remaining fly powder was washed off the spatula with an additional 1ml lysis buffer. Tubes were incubated on ice for 15-30 min, with mixing every 5 min. Epitope-tagged proteins were immunoprecipitated with either anti-Flag- or anti-HA-coated magnetic beads (Sigma and Pierce, respectively, M8823 and 88836) following the manufacturer’s instructions. RNA was eluted from the beads with Proteinase K (Sigma, AM2546) and TRIzol LS Reagent (Ambion, 10296028) was used to isolate immunoprecipitated RNA. The Ovation SoLo RNA-Seq System (Tecan Genomics, S02240) was used to make PABP library. eCLIP libraries were generated by following the protocol in Nostrand et al., 2016 (Van Nostrand et al., 2016), with a minor modification that 8U of RNAse I (Ambion, AM2295) was used per sample. Purified PABP CLIP libraries were submitted for paired-end 75bp sequencing on the Illumina NextSeq platform at the Center for Genomics and Bioinformatics (CGB) in Indiana University, Bloomington. eCLIP libraries were submitted for paired-end 75bp sequencing on the Illumina NextSeq platform at the CGB in Indiana University, Bloomington or paired-end 100bp sequencing on the DNBseq platform at the BGI Genomics in China.

### RNA-seq and PABP CLIP-seq data analysis

RNA-seq datasets were obtained from the following references and are referred to in the text by the surname of the first author: (Doupe et al., 2018; Dutta et al., 2015; Fast et al., 2020; Korzelius et al., 2014; Leader et al., 2018; Li et al., 2018). Both RNA-seq and PABP CLIP-seq read files were processed and aligned to the Berkeley *Drosophila* Genome Project (BDGP) assembly release 6.28 (Ensembl release 99) reference genome using a python based in house pipeline (https://github.com/jkkbuddika/RNA-Seq-Data-Analyzer v1.0). Briefly, the quality of raw sequencing files was assessed using FastQC (Andrews, 2010) version 0.11.9, low quality reads were removed using Cutadapt (Martin, 2011) version 2.9, reads aligning to rRNAs were removed using TagDust2 (Lassmann, 2015) version 2.2, remaining reads were mapped to the Berkeley *Drosophila* Genome Project (BDGP) assembly release 6.28 (Ensembl release 99) reference genome using STAR (Dobin et al., 2013) genome aligner version 2.7.3a and deduplicated using SAMtools (Li et al., 2009) version 1.10. Subsequently, the aligned reads were counted to the nearest overlapping feature suing the Subread (Liao et al., 2019) version 2.0.0 function *featureCounts*. Finally, bigWig files representing RNA-seq coverage were generated using deepTools (Ramirez et al., 2016) version 3.4.2 with the settings --normalizeUsing CPM --binSize 1. All programs, versions and dependencies required to execute the RNA-seq data analyzer are described in the user guide and can be installed using miniconda. The TMM-normalized expression values were computed using the Bioconductor package edgeR (Robinson et al., 2010) version 3.28.1. Genes with TMM-normalized expression value greater than 10 in all replicates were considered as a moderately expressed gene during transcriptome comparisons. Differential gene expression analysis was performed with the Bioconductor package DESeq2 (Love et al., 2014) version 1.26.0 using FDR < 0.05 and log_2_ fold change > 1, unless otherwise noted. For both edgeR-based transcriptome analysis and DESeq2-based differential gene expression analysis only protein coding genes were used. All data visualization steps were performed using custom scripts written in R.

### Enhanced-CLIP (eCLIP) data analysis

The eCLIP read file processing, alignment and processing was done using an easy to use python based in house analysis pipeline, eCLIP data analyzer (https://github.com/jkkbuddika/eCLIP-Data-Analyzer). While the eCLIP data analyzer pipeline permits analysis in both paired and single-end modes, we performed all our analyzes using the single-end mode, which uses only R2 read for analysis. The eCLIP data analyzer pipeline performs the following steps: (1) read quality assessment using FastQC (Andrews, 2010), (2) trimming of universal eCLIP adaptors using Cutadapt (Martin, 2011), (3) addition of UMI sequence (5’-NNNNNNNNNN in the R2 read) using UMI-tools (Smith et al., 2017) to read name to facilitate PCR duplicate removal, (4) removal of reads aligning to rRNAs using Tagdust2 (Lassmann, 2015), (5) alignment of remaining reads to the provided genome (i.e., BDGP assembly release 6.28, Ensembl release 100) using STAR (Dobin et al., 2013), (6) coordinating sort and index alignment outputs using SAMtools (Li et al., 2009), (7) removal of PCR duplicates using UMI-tools (Smith et al., 2017), (8) generation of bigwig files representing RNA-seq coverage tracks using deepTools (Ramirez et al., 2016), (9) quantification of nearest overlapping features using Subread (Liao et al., 2019) function *featureCounts*, and (10) peak calling using PureCLIP (Krakau et al., 2017). All programs, versions and dependencies required to execute the eCLIP data analyzer are described in the user guide and can be installed using miniconda. Input normalized peak calling output files were processed as described in (Busch et al., 2020). Subsequently, significantly enriched *de novo* binding motifs were identified using DREME (Bailey, 2011). Protein coding FMRP target genes were used to identify enriched Gene Ontology (GO) terms using gProfiler. A selected significantly enriched GO categories were plotted using R. Unless otherwise noted, all data visualization steps were performed using custom scripts written in R.

### CLIP-qPCR

Intestine were dissected from 100 *w^1118^* females, and FMRP was precipitated with anti-FMRP antibody (5A11, DSHB) using the same method described above. RNA was isolated from either the starting intestinal lysate or from the IP with Trizol LS (Ambion, 10296028). Resulting RNA was treated with Turbo DNase(ThermoFisher, AM2239) and ~200ng of input RNA and all the RNA derived from IP sample was used for cDNA synthesis with Superscript III (ThermoFisher, 56575). qPCR was performed using the PowerUp SYBR Green Master Mix (ThermoFisher, A25742) on a StepOnePlus machine (ThermoFisher). Primers for all targets detected are listed in Table S5. Transcript levels were quantified in duplicate and normalized to *CG10116*. Fold enrichment was calculated as the ratio of transcript in IP versus input.

## Statistical analysis

Statistical analyses were performed using Prism (GraphPad, Version 7.0). Datasets were tested for normality using D’Agostino-Pearson test. Comparisons of two datasets was performed with an unpaired t-test while qPCR samples were analyzed with a paired t-test. The statistical significance of overlaps in Venn Diagrams was calculated using a hypergeometric test on R. When multiple pairwise comparisons were needed, the R package GeneOverlap was used to perform Fischer’s exact test which yields the statistical significance, odds ratio and Jaccard indices for each pairwise comparison. Significance is indicated as follows: n.s., not significant; *p < 0.05; **p < 0.01; ***p < 0.001; ****p < 0.0001.

## Data availability

The PABP CLIP-seq and FMRP eCLIP-seq datasets from this study have been submitted to the NCBI Gene Expression Omnibus under accession number XXXX (to be added later).

## Acknowledgements

We thank Steven Hou, Gerald Rubin, Steve Stowers, the Kyoto Drosophila Stock Center, the Bloomington Drosophila Stock Center (supported by grant NIHP4OOD018537), the Drosophila Genome Resource Center (supported by grant NIH2P40OD010949), and the Developmental Studies Hybridoma Bank for reagents; the Light Microscopy Imaging Center (supported by grant NIH1S10OD024988-01) for access to the SP8 confocal; and the National Institute of General Medical Sciences (Award R01GM124220) for financial support. The authors declare no competing financial interests.

## Competing Interests

No competing interest declared.

## Supplementary Figure Legends

**Figure S1:**
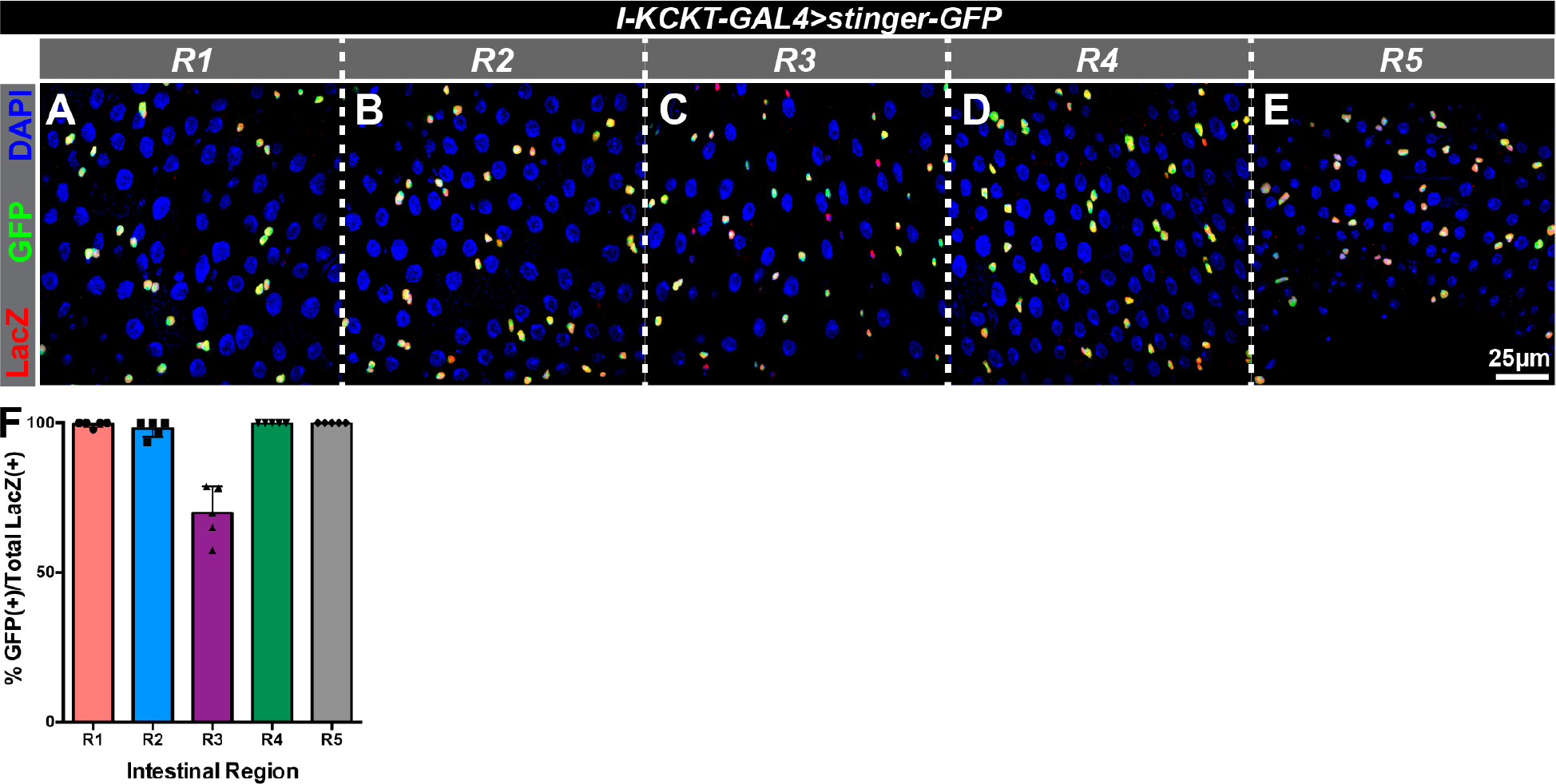
*I-KCKT-GAL4* and *esg-LacZ* label the same cells. (A-E) Micrographs of five intestinal regions (R1-R5) stained for *I-KCKT-GAL4*-driven *stinger-GFP* in green, the intestinal progenitor marker *esg-LacZ* in red, and the DAPI DNA marker in blue. (F) Histogram of the percentage of *esg-LacZ*-positive cells that also *-stinger-GFP* in each region.

**Figure S2:**
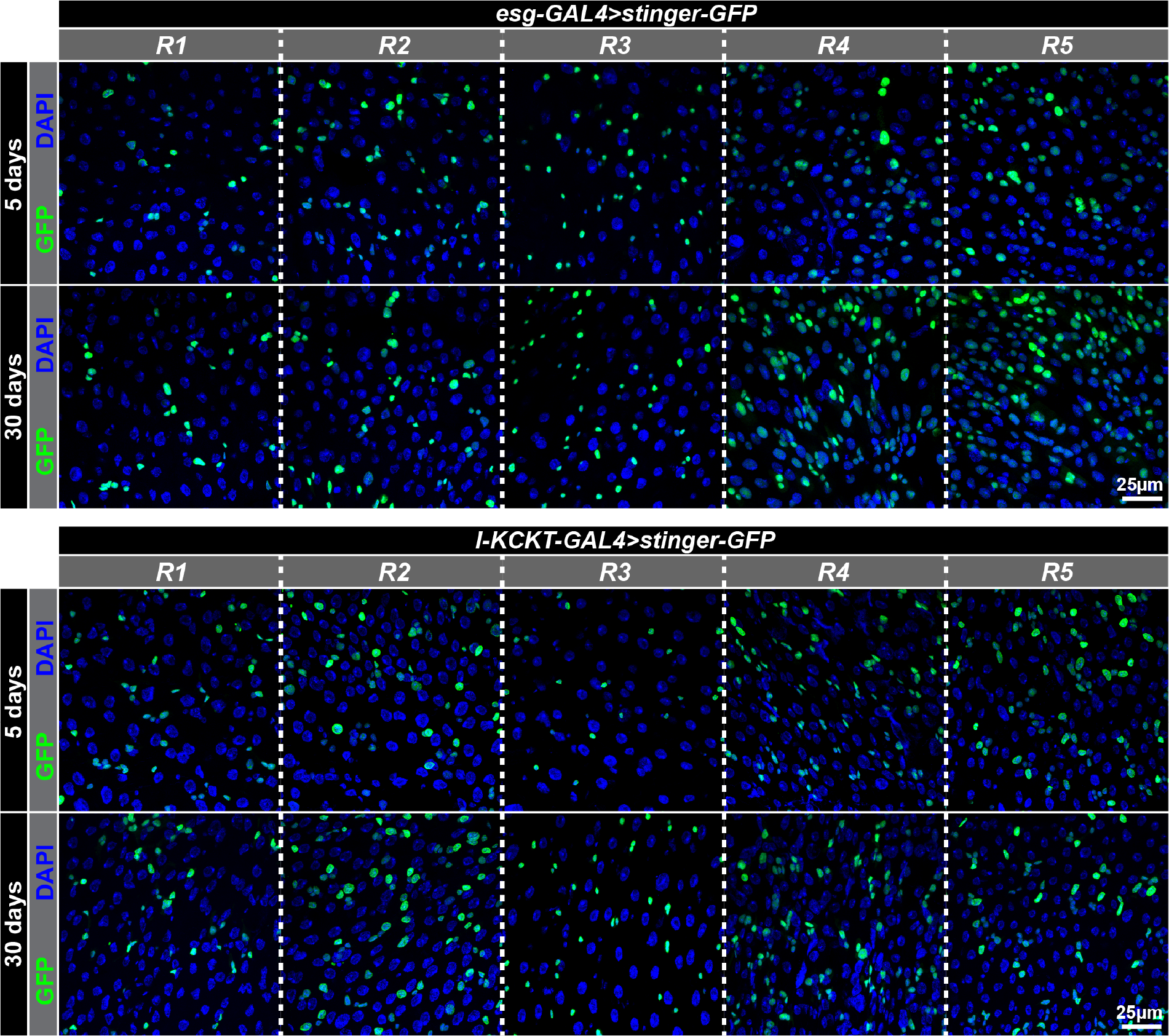
Comparison of *I-KCKT-GAL4* and *esg-GAL4* expression in aged flies. Representative micrographs of the five intestinal regions (R1-R5) of young (5 day) and older (30-day) females showing *UAS-stinger-GFP* driven by either *esg-GAL4* (top) or *I-KCKT-GAL4.p65* (bottom) and counterstained for the DAPI DNA marker in blue.

**Figure S3:**
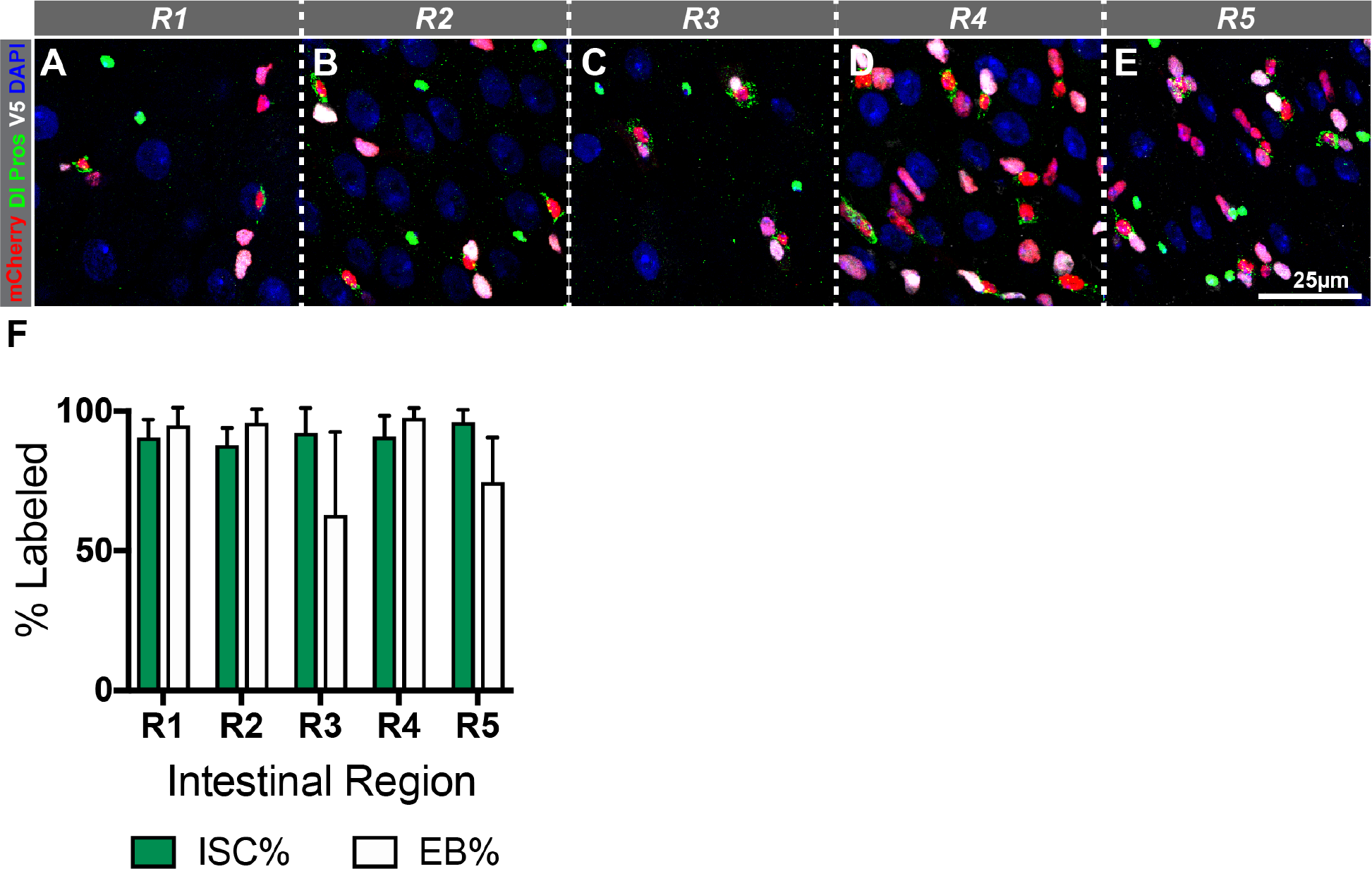
Verification that dual reporters *mira-His2A.mCherry.HA* and *3Xgbe-smGFP.V5.nls* distinguish ISCs and EBs. (A-E) Representative micrographs of the five regions (R1-R5) from the intestines of 5-day old females stained for ISCs (anti-Delta, green cytoplasmic staining), EEs (anti-Prospero, green nuclear staining), *mira-His2A.mCherry.HA* (red), *3Xgbe-smGFP.V5.nls* (*white*), and the DAPI DNA marker (blue). (F) Quantification of the percent of ISCs and EBs labeled by the dual reporters *mira-His2A.mCherry.HA* and *3Xgbe-smGFP.V5.nls* in each of the five intestinal regions. ISCs were defined as the number of Delta+ cells per field of view. EBs were defined as the number of Delta-, mCherry+ cells per field of view. For quantification, field of view images were taken and quantified from defined portions of each intestinal region in 6-7 intestines. We note that we detected a few (1-5) cells that were both Delta+ and *3Xgbe-smGFP.V5.nls+* in regions of most intestines. These were not scored as either ISCs or EBs, and could represent cells transitioning between these two cell fates.

**Figure S4:**
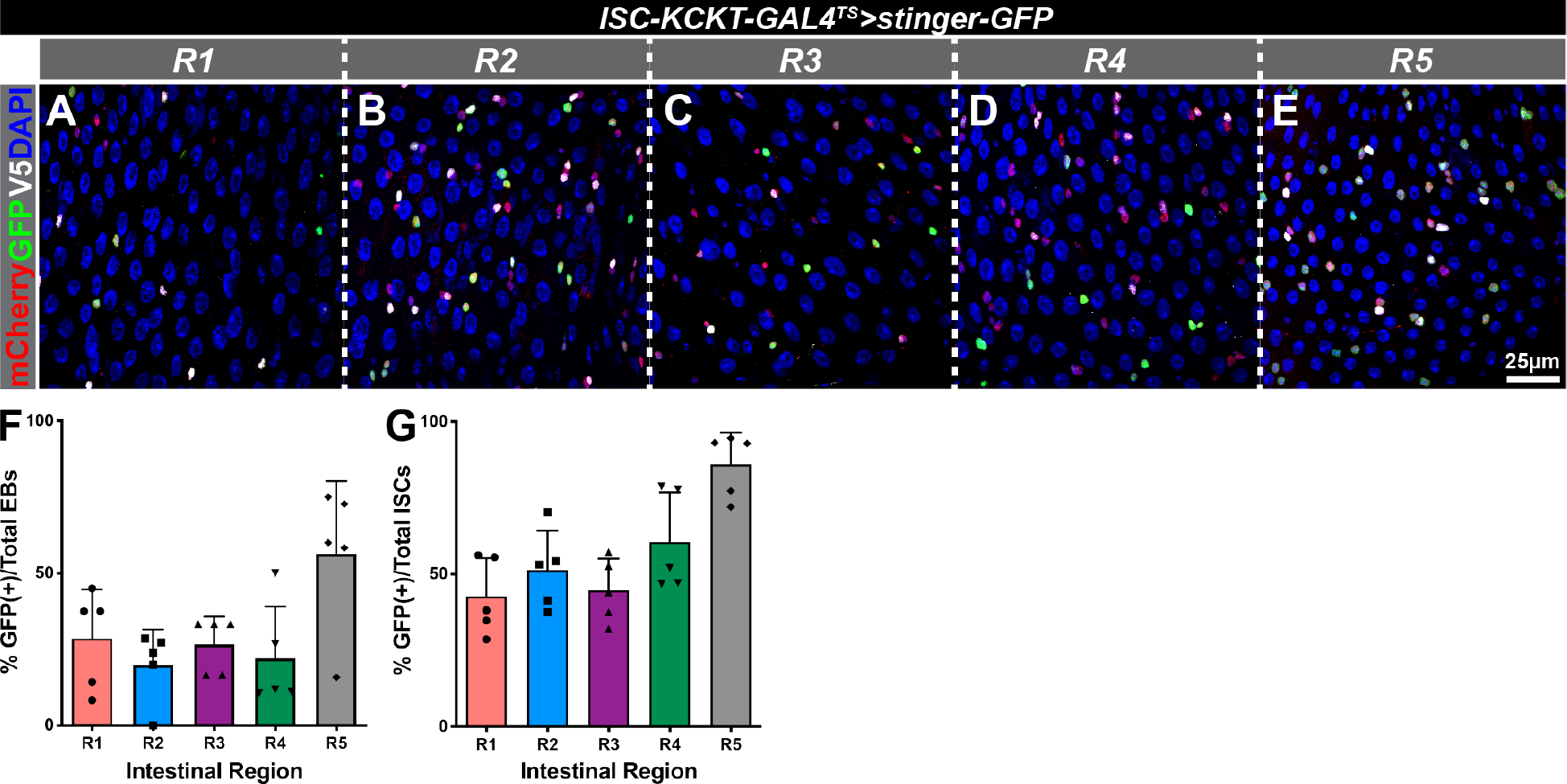
*ISC-KCKT-GAL4^TS^* labels many ISCs. (A-E) Representative micrographs of the five regions (R1-R5) from the intestines of 5-day old females stained for *ISC-KCKT-GAL4^TS^*-driven *UAS-stinger-GFP* in green, *mira-His2A.mCherry.HA* in red, *3Xgbe-smGFP.V5.nls* in white, and the DAPI DNA marker in blue. ISCs were scored as cells positive for *mira-His2A.mCherry.HA* and negative for *3Xgbe-smGFP.V5.nls*, while EBs were scored as cells positive for both markers. (F) Histogram of the percentage of EBs that expressed *ISC-KCKT-GAL4^TS^-driven UAS-stinger-GFP* in each region. (G) Histogram of the percentage of ISCs that expressed *ISC-KCKT-GAL4^TS^*-driven *UAS-stinger-GFP* in each region.

**Figure S5:**
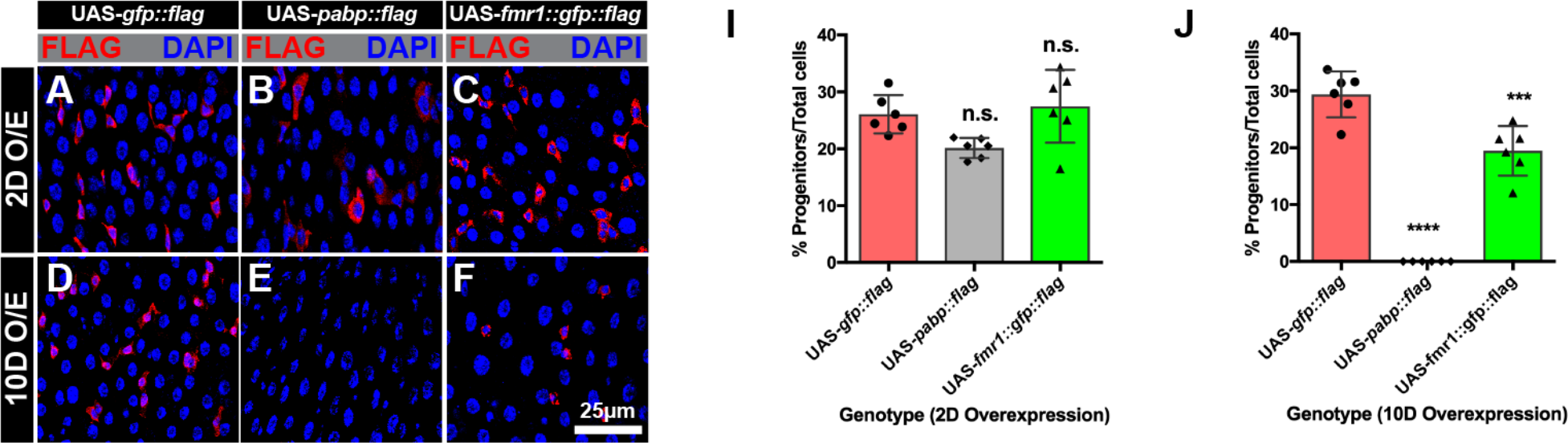
*I-KCKT-GAL4^TS^-dependent* expression of tagged RBPs in intestinal progenitor cells. (A-J) Micrographs of intestines showing *I-KCKT-GAL4^TS^-dependent* expression of *UAS-GFP.FLAG* (A, D), *UAS-PABP.FLAG* (B,E), and *UAS-FMR1.GFP.FLAG* (C,F) after two (A-C) and ten (D-F) days at the permissive temperature. Expressed proteins were detected with either anti-FLAG or -HA antibodies (shown in red) and intestines were counterstained with the DAPI DNA marker in blue. Full genotypes are listed in Table S4. (I, J) Histograms of the percentage of progenitor cells after two and ten days of *I-KCKT-GAL4^TS^-dependent* expression, based on quantification of the numbers of HA+ or FLAG^+^ and DAPI^+^ cells.

**Figure S6:**
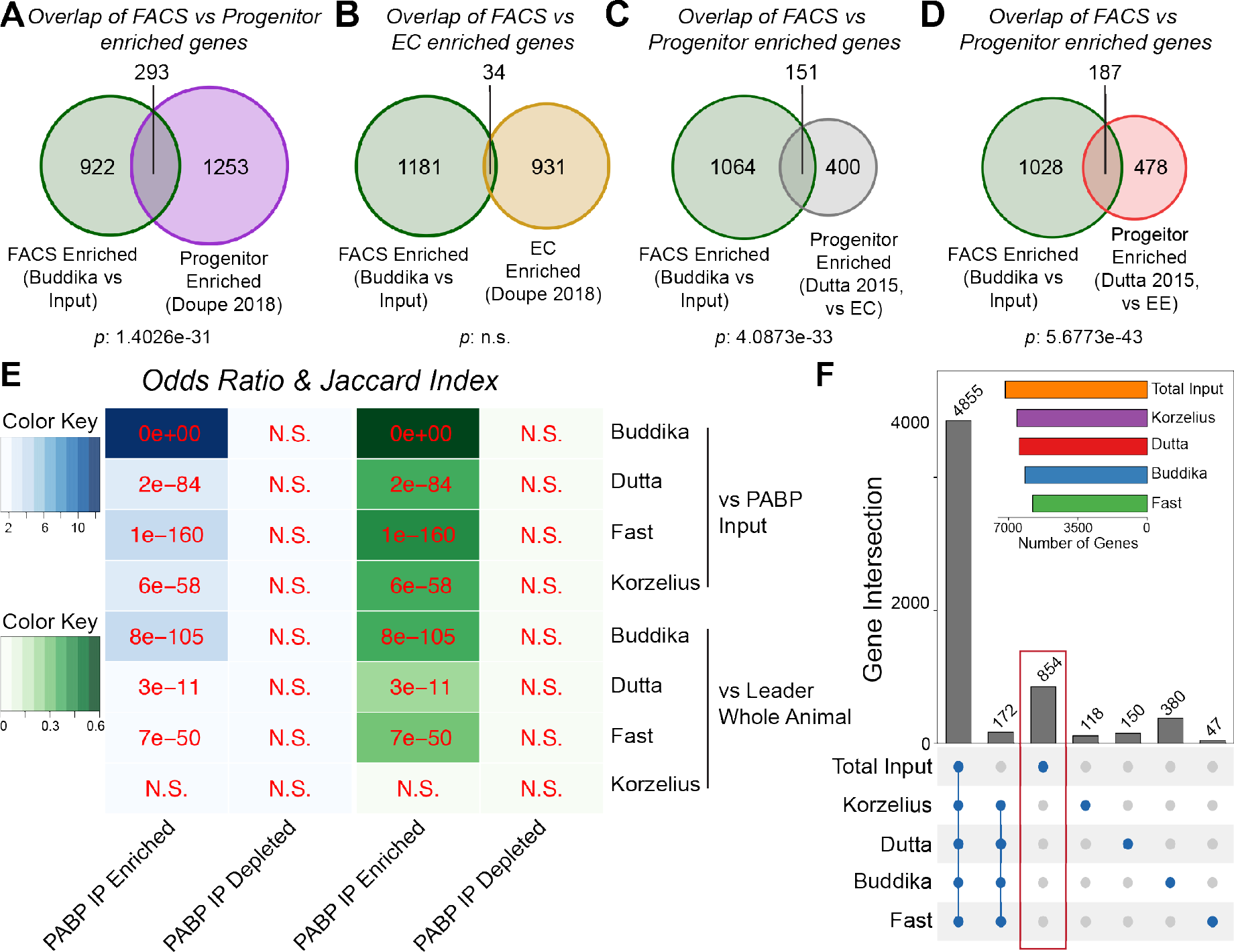
I-KCKT based RIPs identifies genes expressed in intestinal progenitors. (A-D) Venn diagrams showing genes significantly enriched in FACS-isolated intestinal progenitor data from Buddika that overlap with Doupe’s progenitor enriched genes (A), Doupe’s EC enriched genes (B), Dutta’s progenitor enriched genes as compared to EC genes (C), or Dutta’s progenitor enriched genes as compared to EE genes (D). (E) Heatmaps of odds ratio (in blue) and Jaccard index (in green) values for pairwise comparisons of PABP IP enriched or PABP IP depleted genes against input normalized FACS-based progenitor RNA-seq. Two sources of input data have been used for FACS data normalization: total input of the PABP experiment, or the female whole animal dataset from Leader. Numbers on each colored box show the p-value for each overlap based on Fisher’s exact test. N.S.: Not significant. (F) Upset plot showing the overlap of moderately expressed genes identified by FACS based methods and total PABP IP input. Only a select set of meaningful overlaps are shown. The number above each bar shows the precise size of each intersection. Note that 854 genes were uniquely present in the input, but not in any of the FACS based datasets

**Figure S7:**
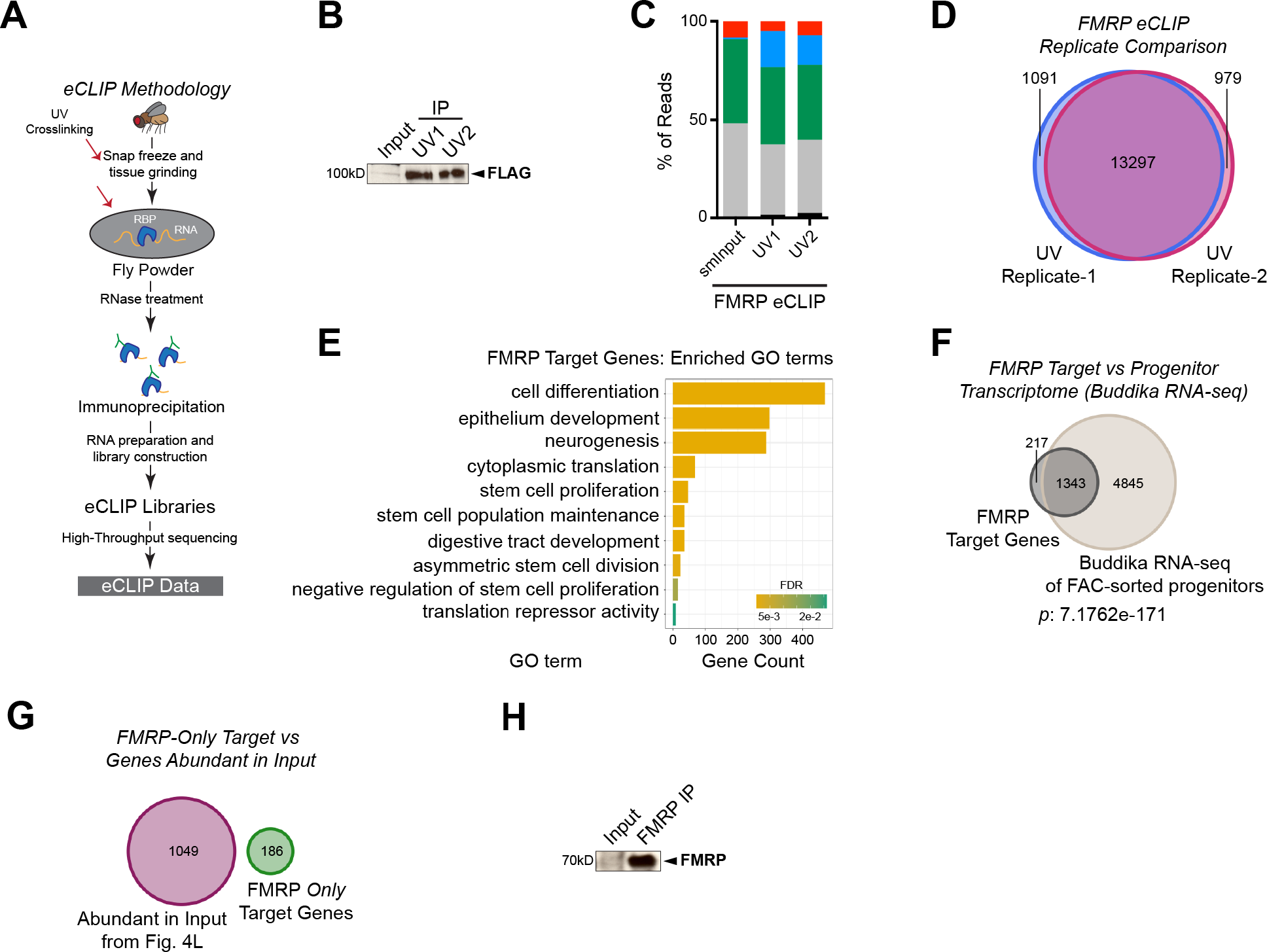
FMRP eCLIP using *I-KCKT* identifies target transcripts. (A) Schematic representation of the eCLIP methodology. (B) Western blot of input and FLAG IP samples (UV1 and UV2) prepared from whole animal extract and probed with anti-FLAG antibodies. (C) Grouped bar plot showing the percentage of sequencing read distribution. (D) Venn diagram showing the overlap of eCLIP peaks identified in UV replicate 1 and 2. (E) Bar plot showing a selected set of significantly enriched biological process Gene Ontology (GO) terms for FMRP target genes. (F) Venn diagram of FMRP target genes and intestinal progenitor transcriptome identified by RNA-seq of Buddika. (G) Venn diagram of top 10% genes with highest TMM normalized expression values from Fig. 4L and FMRP target genes that are absent in the progenitor transcriptome (186 genes from Fig. 5D). Note that the 186 genes are unlikely to be contaminants as they do not overlap with most highly abundant genes in the preparation. (H) Western blot of input and FMRP IP samples prepared from dissected intestines probed with anti-FMRP antibodies.

## Supplementary Tables

Descriptions of the five supplementary Tables, supplied as excel files, are included below.

Table S1: List of genes differentially expressed in PABP IP versus whole animal Input.

Table S2: List of genes common to either PABP IP or FACS-isolated progenitors (Buddika 2020b) and DamID-identified progenitor or EC specific genes (Doupe et al., 2018).

Table S3: List of genes genes identified by FMRP eCLIP.

Table S4. List of fly genotypes in figures.

Table S5. List of DNA primers.

## Notes

### Competing Interest Statement

The authors have declared no competing interest.

### Summary of Updates

Added Figure 5; Results, Discussion, and Method updated.

